# The importance of input sequence set to consensus-derived proteins and their relationship to reconstructed ancestral proteins

**DOI:** 10.1101/2023.06.29.547063

**Authors:** Charlotte Nixon, Shion A. Lim, Matt Sternke, Doug Barrick, Mike Harms, Susan Marqusee

## Abstract

A protein sequence encodes its energy landscape - all the accessible conformations, energetics, and dynamics. The evolutionary relationship between sequence and landscape can be probed phylogenetically by compiling a multiple sequence alignment of homologous sequences and generating common ancestors via Ancestral Sequence Reconstruction or a consensus protein containing the most common amino acid at each position. Both ancestral and consensus proteins are often more stable than their extant homologs - questioning the differences and suggesting that both approaches serve as general methods to engineer thermostability. We used the Ribonuclease H family to compare these approaches and evaluate how the evolutionary relationship of the input sequences affects the properties of the resulting consensus protein. While the overall consensus protein is structured and active, it neither shows properties of a well-folded protein nor has enhanced stability. In contrast, the consensus protein derived from a phylogenetically-restricted region is significantly more stable and cooperatively folded, suggesting that cooperativity may be encoded by different mechanisms in separate clades and lost when too many diverse clades are combined to generate a consensus protein. To explore this, we compared pairwise covariance scores using a Potts formalism as well as higher-order couplings using singular value decomposition (SVD). We find the SVD coordinates of a stable consensus sequence are close to coordinates of the analogous ancestor sequence and its descendants, whereas the unstable consensus sequences are outliers in SVD space.

## Introduction

With the flood of DNA sequences available for many protein families, we now have access to protein sequences for thousands of different homologs. These sequences provide valuable information for protein engineering studies. Computational analyses of multiple sequence alignments (MSAs) can be used to generate novel sequences that encode proteins with novel properties^1–5^. Two common approaches for this are ancestral sequence reconstruction (ASR) and consensus design. ASR takes advantage of the evolutionary relationships among homologs, which are used to reconstruct sequences of ancestral proteins along the phylogenetic tree^6, 7^. Alternatively, these same MSAs can be used to identify consensus residues as the most common residue at each position in the MSA, independent of their sequence context^2^. These consensus residues can be combined to give a *de novo* protein sequence comprising the full consensus or, alternatively, to modify select residues in an extant sequence, resulting in a variant with novel, desired properties. Both approaches have been successfully used to generate new proteins that retain function and often show increased thermodynamic stability^8–11^. Highly conserved residues are likely to be selected in both approaches, and have led some to suggest that ASR does little more than provide a bias towards consensus residues^12–14^. Here, we have used the Ribonuclease H1 (RNase H) family to compare these two approaches and evaluate how the evolutionary relationship of the sequences in the input set affects the resulting consensus protein. We address two important questions: 1) How does a consensus protein compare to an ancestral protein? and 2) How do the biophysical properties of a consensus protein depend on the sequences used for the MSA to generate the consensus (both the size of the input set and their evolutionary relationship)?

RNase H is a small, single domain protein that degrades RNA strands of RNA/DNA hybrids.^15^ The energetics and folding pathway of both *E. coli* RNase H (ecRNH) and *T. thermophilus* RNase H (ttRNH) have been studied extensively^16–22^. Based on this wealth of biophysical data and the availability of hundreds of homologs, the evolutionary divergence of thermostability along the mesophilic and thermophilic lineages was characterized by ASR^23, 24^. Anc1 represents the node connecting the mesophilic and thermophilic branch and is the last common ancestor connecting the two extant RNases H: ecRNH, a mesophilic protein with a T_m_ of 68 °C, and ttRNH, a thermophilic protein with a T_m_ of 88 °C^19^. From Anc1, thermostability was found to diverge along the two lineages, with T_m_ increasing along the thermophilic lineage and decreasing along the mesophilic lineage. Notably, this common ancestor (Anc1) did not exhibit the hyperstability commonly seen in ASR studies on other protein families. Because of this, and the fact that consensus proteins are often thought to generate hyperstable proteins, we investigated the extent and origin of the differences between ancestral and consensus proteins in the RNase H family.

We generated consensus proteins using different input sequences derived from different lineages of the RNase H tree and compared their biophysical properties with the previously studied ancestral proteins. The consensus protein derived using members from the whole phylogenetic tree (the same 405 sequences used in the previous ASR study) is an active RNase H protein, yet it has lower thermodynamic stability and displays a less cooperative folding transition than ancestral and extant RNases H including the common ancestor Anc1. When input sequences are restricted to those more closely related to ecRNH, however, the resulting consensus protein is notably stabilized, comparable to many previously studied extant and ancestral RNases H. These findings show that the input set of sequences can have a large effect on the resulting energetic properties of a consensus protein, and that consensus proteins derived from a phylogenetically diverse MSA can produce relatively unstable proteins.

To tease apart the origins of these differences, we used an evolutionary coupling analysis to compare the consensus and ancestral sequences. Both the consensus and ancestral proteins retain similar co-evolutionary total coupling scores, but differed in the details of the co-evolutionary signal. When we compared the coupling patterns in each sequence using singular value decomposition, the extant and ancestral sequences formed clusters with one another. In contrast, only consensus sequences calculated for phylogenetically restricted subsets of the alignments were part of these clusters; consensus sequences calculated from the entire alignment did not. Finally, our results demonstrate that starting with the same MSA, ASR and consensus approaches yield proteins with very different properties, which would not be expected if the two methods provided different routes to the same endpoint.

## Results

### Generation of Consensus RNases H

We generated a consensus bacterial RNase H based on the 405 sequences and MSA used in the previous ASR studies^23^. The resulting consensus protein, named WholeCons* (asterisk indicates cysteine-free variant), encodes the most common residue at each position of the MSA (Figure 1A). WholeCons* shares 78% sequence identity with ecRNH*, 64% with ttRNH*, and 76% with the common ancestor Anc1*. Given the large number of bacterial RNase H sequences now available, we also generated a consensus RNase H using a larger alignment of 3,312 RNases H sequences, named MegaCons*. Despite the eightfold increase in the number of sequences, MegaCons* differs from WholeCons* at only eight positions indicating that the ASR input sequence set (n = 405) reasonably captures the sequence diversity of the RNase H family.

**Figure 1:**
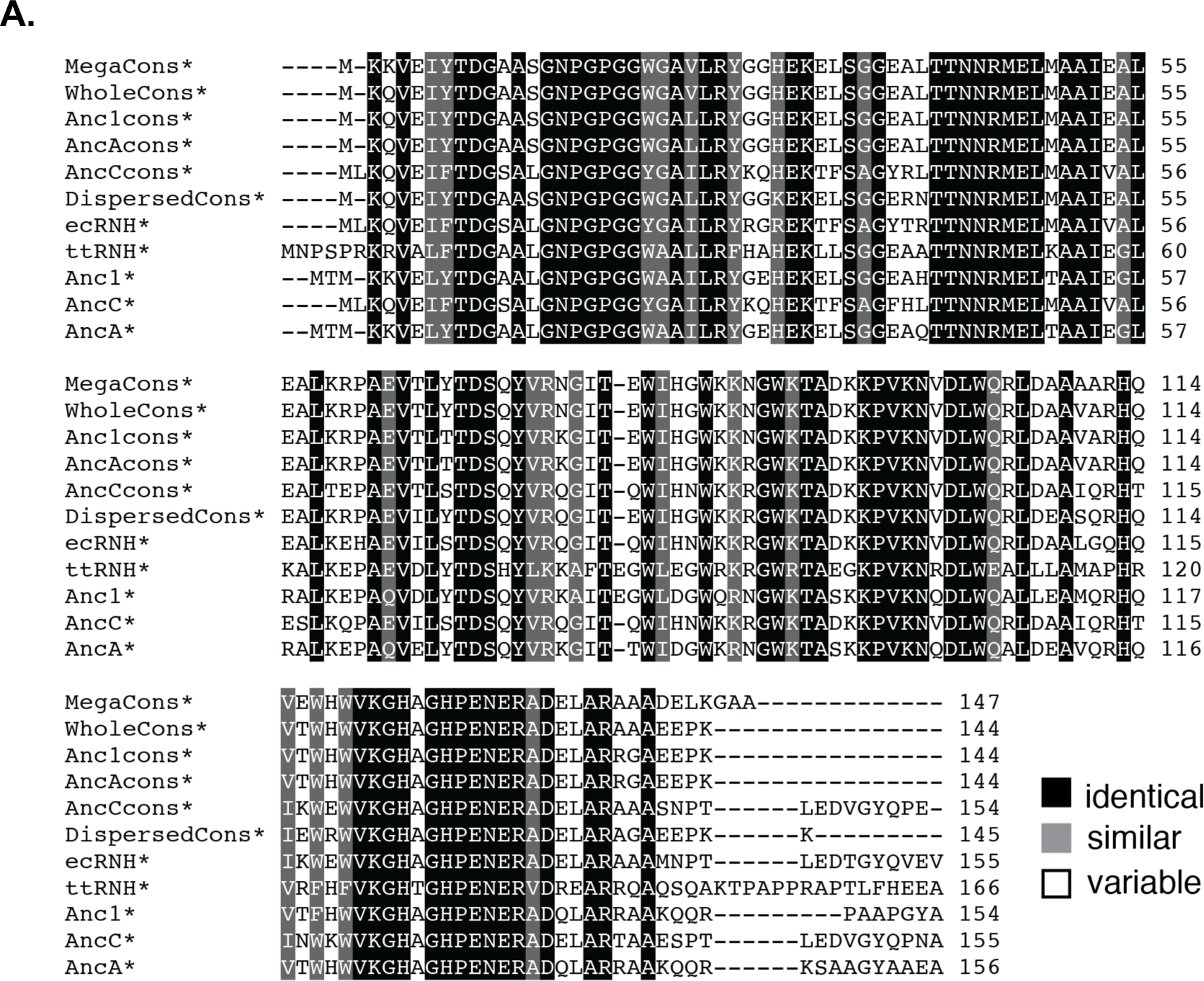
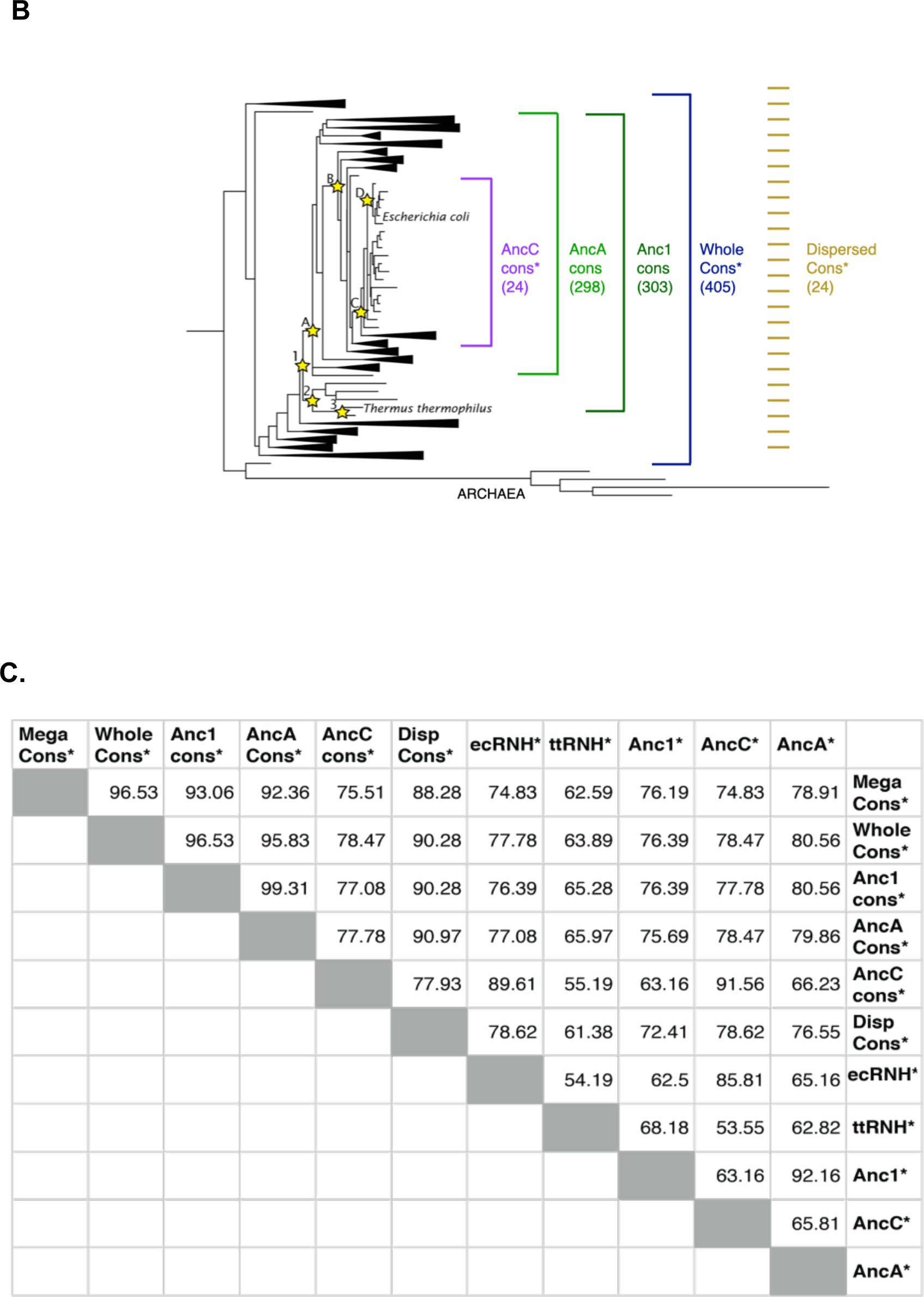
Sequence comparisons of extant, ancestors and consensus RNases H. (**A).** MSA of the different consensus, ancestral^23^, and extant RNases H^19^. Alignments were generated with Clustal Omega^52^ and shaded by similarity^53^. (**B).** RNase H phylogenetic tree with characterized ancestors highlighted as stars.^23^ Designed consensus proteins are displayed as brackets showing the input extant sequence sets, with the number of input sequences in parentheses. **(C).** Percent identity calculated by Clustal Omega^52^ between consensus and selected ancestral^23^ and extant^19^ RNases H. (DispersedCons* abbreviated DispCons*).

To explore how the phylogenetic diversity of the input sequence set affects the sequence and biophysical properties of the resulting consensus protein, we generated alternative consensus proteins using different phylogenetically restricted subsets of the 405 sequences used in the tree (Figure 1B). Anc1cons is the consensus of homologs that descend from Anc1 (303 sequences). AncAcons is the consensus of homologs that descend from AncA, the first ancestor in the mesophilic lineage (298 sequences). We did not generate a consensus from the Anc2 (the first ancestor in the thermophilic lineage) descendants due to the small number of input sequences (5 sequences). Additionally, we generated AncCcons* using the 24 homologs descended from AncC, which represents a consensus protein designed from a set of closely related sequences. To evaluate the relative effects of taxonomic restriction versus sequence reduction in AncC, we also generated DispersedCons*. DispersedCons*, like AncCcons*, is also derived from only 24 sequences, however in this case we used distantly related sequences chosen at a regular interval in the alignment.

The different consensus proteins display a broad range of sequence identity (55 to 92%) to the extant and ancestral sequences (Figure 1C). Figure 1A shows an alignment of the consensus sequences with select extant and ancestral RNases H, which demonstrate the dependence of the consensus approach on the input sequences. Clearly, it is not possible to define a single consensus sequence for the RNase H family without defining the input sequences.

### WholeCons* folds into an active RNase H with low apparent stability and cooperativity

WholeCons* was expressed and purified from *E. coli* cells using established RNase H purification protocols (see Methods). The protein is enzymatically active and its circular dichroism (CD) spectrum is consistent with those previously observed for the RNase H fold^23^ (Figure 2). The equilibrium stability of WholeCons* was probed by thermal and chemical denaturation monitoring the CD signal at 222 nm (Figure 3A,B) and analyzed using a two-state model^25^. The thermal unfolding curve is reversible and results in a T_m_ of 62.8 ± 0.2 °C, notably lower than that for ecRNH and Anc1 (68.0 ± 0.5 °C. and 88.5 °C, respectively). Likewise, the urea-induced unfolding transition occurs at a lower urea concentration than that of ecRNH and Anc1; the two-state linear extrapolation analysis results in a ΔG°_apparent_ (25 °C) of 5.1 ± 0.1 kcal/mol (Table 1), significantly lower than the range of 9.7 to 12.8 kcal/mol observed for ecRNH, ttRNH, and the ancestral RNases H. The fitted *m*-value of 1.5 ± 0.1 kcal/mol M^−1^, is also notably lower than typically observed for RNase H (∼1.9 – 2.1 kcal/mol M^−1^), suggesting either that urea-induced unfolding is multistate, or that the native state of WholeCons* is partially disordered.

**Figure 2:**
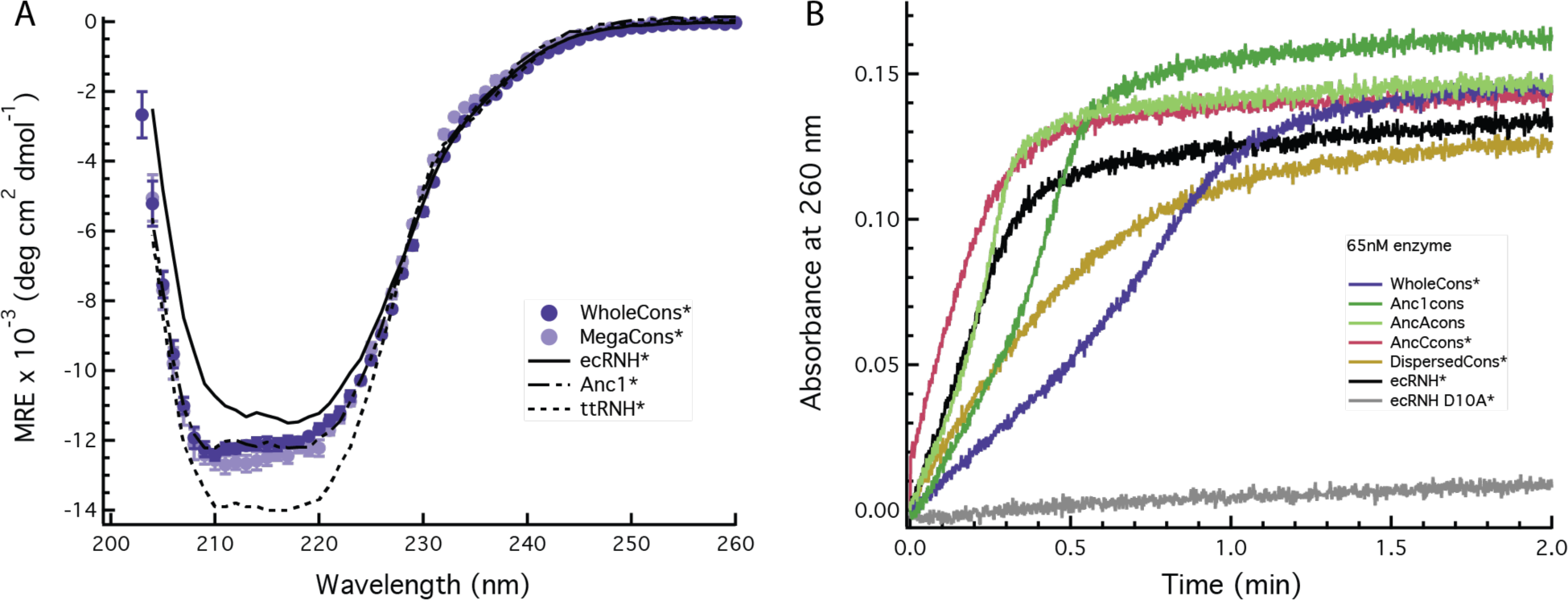
Consensus proteins adopt the RNase H fold and are active. **(A).** CD spectra and **(B).** RNase H activity assays for consensus RNase H constructs compared to extant and ancestral proteins.^24^ ecRNH D10A* is a catalytically-inactive RNase H variant.

**Figure 3:**
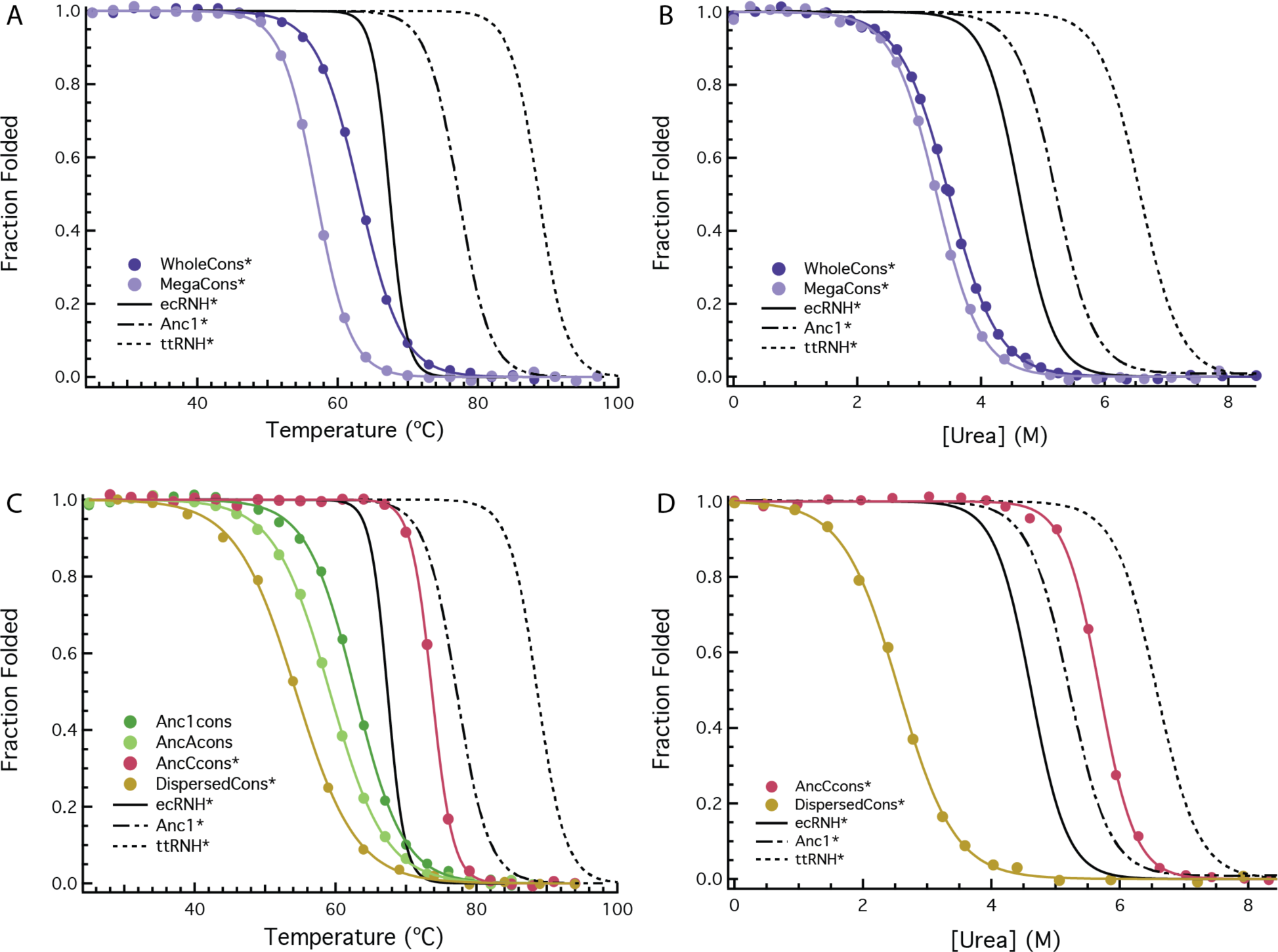
Thermal and chemical denaturation of RNase H variants. **(A)**. Thermal denaturation of WholeCons* and MegaCons* compared to ecRNH, ttRNH, and Anc1^23^. (**B).** Urea denaturation of WholeCons* and MegaCons* compared to ecRNH*, ttRNH*, and Anc1*^24^. (**C).** Thermal denaturation of Anc1cons, AncAcons, AncCcons*, and DispersedCons* compared to ecRNH, ttRNH, and Anc1^23^ and (**D).** Urea denaturation of AncCcons*, and DispersedCons* compared to ecRNH*, ttRNH*, and Anc1*^24^. All unfolding transitions are monitored by CD spectroscopy at 222 nm.

**Table 1:** Thermodynamic parameters for temperature and urea-induced unfolding assuming a two-state linear free energy model^25^ for consensus proteins and core fragments compared to extant^18, 21^ and ancestral^24^ proteins. (pH 5.5 and ΔG_unf_ and *m*-values at 25 °C).

| | T<sub>m</sub> (°C) | $\Delta G_{\text{unf}}$ (kcal/mol) | $m$-value (kcal/mol/M) | C<sub>m</sub> (M) |
| --- | --- | --- | --- | --- |
| WholeCons* | 62.8 ±0.2 | 5.1 ±0.1 | 1.5 ±0.1 | 3.5 ±0.1 |
| MegaCons* | 56.9 ±0.2 | 5.6 ±0.2 | 1.7 ±0.1 | 3.3 ±0.1 |
| Anc1cons | 61.4 ±1 | - | - | - |
| AncAcons | 58.6 | - | - | - |
| AncCcons* | 76.3 ±2.3 | 11.8 ±0.6 | 2.1 ±0.1 | 5.7 ±0.1 |
| DispersedCons* | 53.3 ±1.0 | 3.5 ±0.1 | 1.4 ±0.1 | 2.6 ±0.1 |
| ecRNH* | 68.0 ±0.5 | 9.7 ±0.4 | 2.1 ±0.09 | 4.62 ±0.04 |
| ttRNH* | 88.5 | 12.8 | 2 | 6.4 |
| Anc1* | 76.7 ±0.3 | 10.2 ±0.2 | 1.91 ±0.03 | 5.34 ±0.06 |
| AncA* | 69.7 ±0.2 | 9.4 ±0.2 | 2.22 ±0.10 | 4.17 ±0.09 |
| AncC* | 67.1 | 9.3 ±0.4 | 2.21 ±0.13 | 4.12 ±0.04 |
| ecRNH* core |  | 3.0 ±0.3 | 1.16 ±0.05 |  |
| ttRNH* core |  | 2.6 ±0.8 | 1.00 ±0.09 |  |
| Anc1*core |  | 1.71 ±0.11 | 0.82 ±0.04 |  |
| WholeCons*core |  | 3.6 ±0.1 | 1.3 ±0.1 |  |

Errors reported are standard deviations of replicate experiments.

To assure that the low apparent stability and *m*-value did not originate from the relatively small number of sequences used to generate WholeCons*, we evaluated the behavior of MegaCons* (derived from 3,312 RNase H sequences). These two consensus sequences are over 96% identical, and the eight amino acid changes result in only minor changes in the thermodynamic properties (Figure 3 and Table 1).

### Lack of cooperativity and two-state behavior in WholeCons*

The low apparent *m*-value points to a potential loss of two-state behavior in the consensus protein. Since *m-*values derived from two-state unfolding transitions correlate with the change in accessible surface area between the unfolded and folded state^26–28^, proteins of the same size and overall composition, such as homologous proteins, are expected to have similar *m*-values. A low *m*-value can arise from either a difference in the change in solvent exposed surface area (via a structural change in the native or unfolded state) or a deviation from the two-state mechanism^29–31^. To test for two-state behavior, we monitored urea-induced denaturation of WholeCons* using two different probes: CD (monitoring secondary structure) and intrinsic tryptophan fluorescence (monitoring packing around the tryptophan). Unlike ecRNH*, where the CD and fluorescence curves overlay within error^16^, the curves for WholeCons* are non-coincident (Figure 4), demonstrating that one or more intermediates are significantly populated in the unfolding transition.

**Figure 4:**
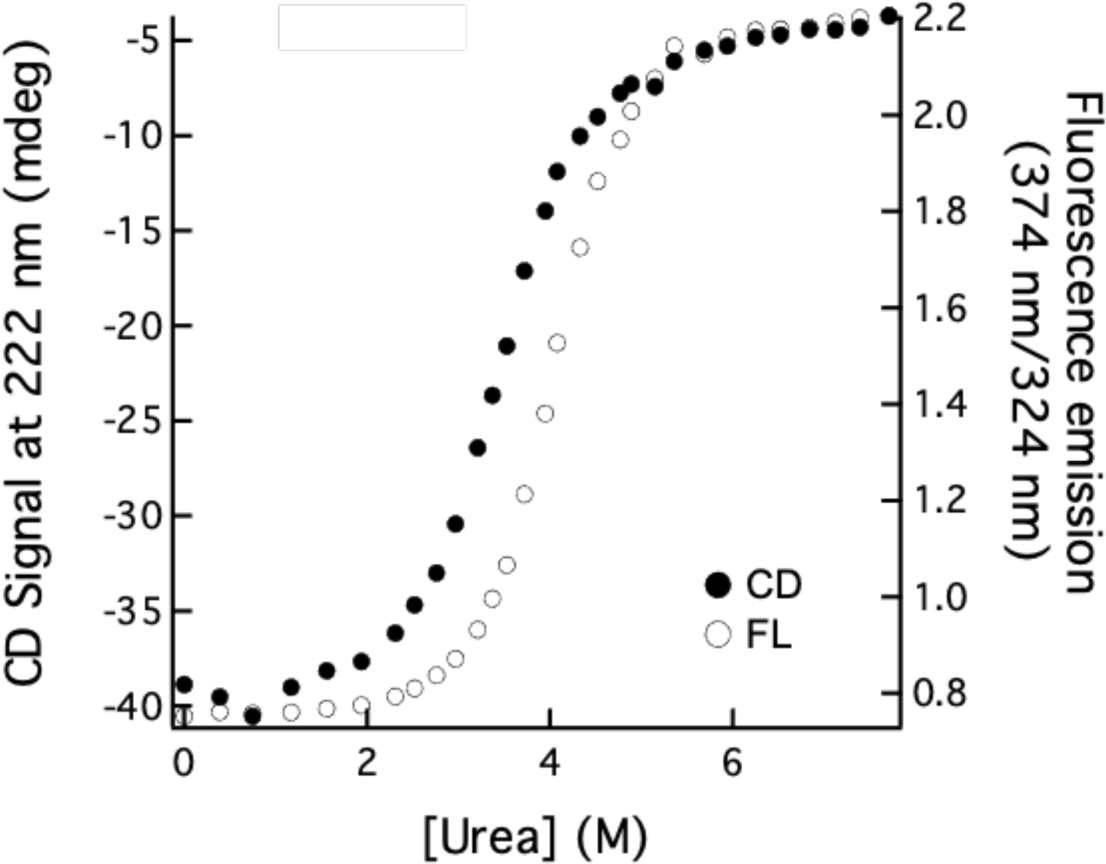
WholeCons* shows non-coincident denaturation curves. CD at 222 nm (filled circles) and tryptophan fluorescence (open circles) (pH 5.5, 25 °C).

Though extant RNases H show two-state equilibrium behavior, all extant RNases H studied to date transiently populate a kinetic intermediate during the refolding process^17, 18, 21^. This canonical RNase H kinetic intermediate forms within the deadtime of the stopped-flow CD instrument (∼10 msec) with about ∼70% of the native CD signal; this burst-phase intermediate converts to the native state by a single exponential on the order of seconds.^17^ Refolding of WholeCons* deviates from this behavior. Under native conditions, the complete signal change for WholeCons* takes place during the burst phase, with no obvious observable phase (Figure 5A). The rapid formation of structure with a lower than expected equilibrium *m*-value suggest that the protein may populate a partly folded intermediate under native conditions without significant population of the fully-folded native state.

**Figure 5:**
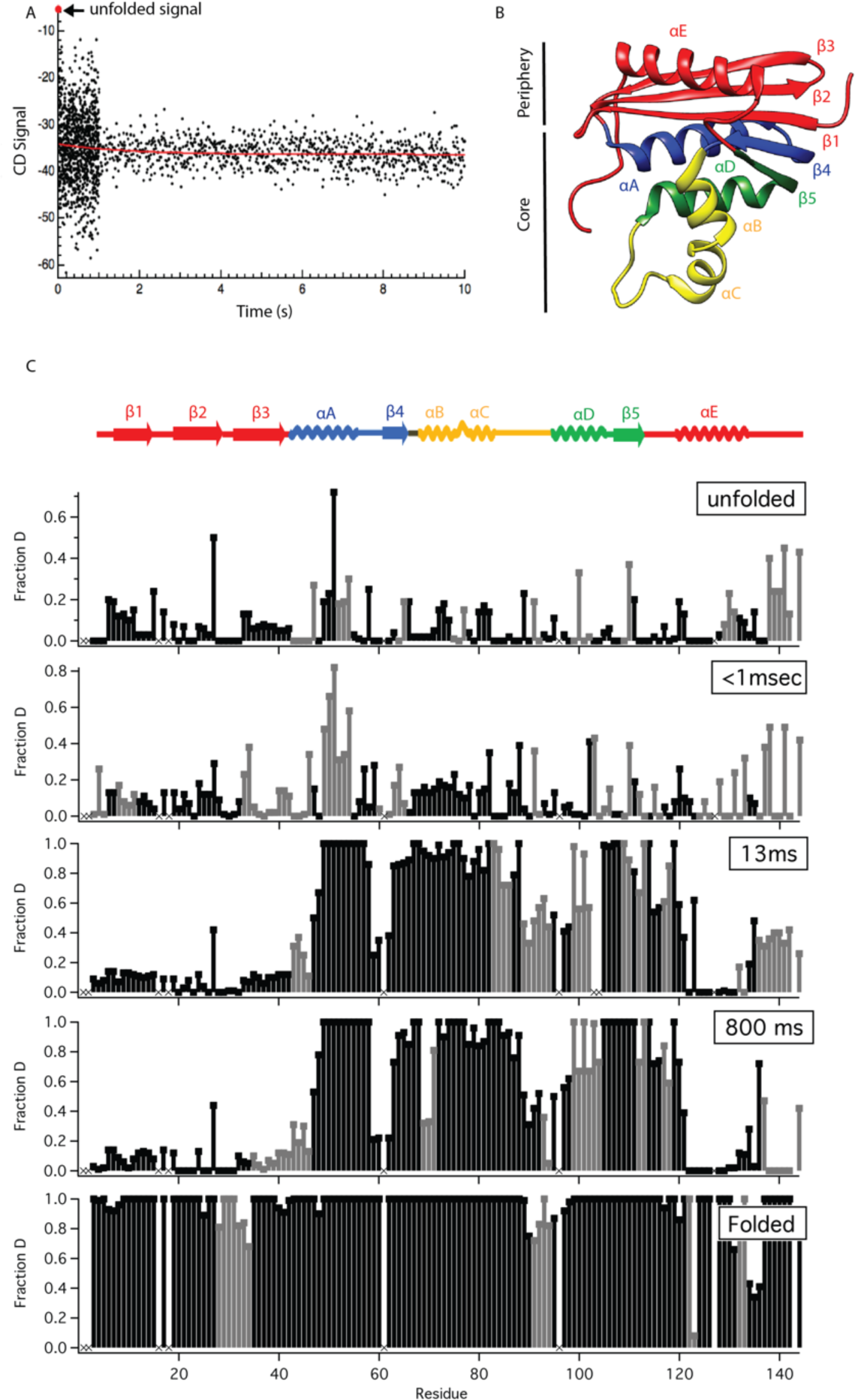
Refolding of WholeCons*. **(A)** Stopped-flow refolding of WholeCons where almost the entire signal change takes place in the dead time. **(B)** *E. coli* RNase H structure (PDB ID: 2RN2^54^) adapted from Hu et al. PNAS. 2013^32^, colored by core (blue, yellow, green) and periphery (red). **(C)** Pulse-labeling hydrogen-deuterium exchange refolding studies monitored by mass spectrometry (HXMS) of WholeCons*. Fraction deuterated is plotted for each residue at different refolding times. Residues in black are site-resolved, residues in grey are not resolved from their neighbors, and residues marked as “x” had insufficient peptide coverage to derive a fraction deuterated. The secondary structural elements of ecRNH core (blue, yellow, green) and periphery (red) are indicated above.

To further characterize the rapid formation of structure seen by WholeCons*, we turned to pulse-labeling hydrogen-deuterium exchange monitored by mass spectrometry^32^. Upon initiation of refolding of WholeCons*, rapid protection is seen in the core region (helices ABCD, strands 4,5; Figure 5 B,C), reminiscent of early folding intermediates seen for extant and ancestral RNases H^32, 33^. However, the peripheral region of WholeCons* remains unprotected at the latest timepoint. The periphery does, however, show protection when subjected to a pulse in the fully folded protein under equilibrium conditions – suggesting observable protection forms on a very slow timescale. It is important to note that protection in these pulsed-experiments requires only minimal residue-level stability (1-2 kcal/mol under the pulse conditions). Taken together, these data are consistent with WholeCons* populating a stable intermediate showing a similar protection pattern (structure) as extant and ancestral RNases H with a very slow and possibly marginal sampling of the native structure. The population of the intermediate under native conditions would account for the low apparent *m*-value seen in WholeCons* equilibrium experiments.

### Folding of a WholeCons* fragment corresponding to the structured regions of the intermediate

Since the pulsed-labeling studies above indicated that the core region of WholeCons* can fold autonomously, we generated a peptide fragment corresponding to this region (residues 41-122, ‘WholeCons* Core’). For many RNase H variants, similar peptide fragments can fold in isolation and serve as an equilibrium mimic of the transient folding intermediate^34^. Indeed, WholeCons* Core folds under native conditions with a CD signal similar to those observed for the core of other RNase H variants; urea denaturation shows a cooperative transition that can be fit with a two-state assumption. The resulting stability and *m*-value (ΔG_unf_(25 °C) = 3.6 ± 0.1 kcal/mol, *m*-value of 1.3 ± 0.1 kcal/mol M^−1^) are comparable to those observed for other RNase H core fragments, which range from 1.3 to 3.4 kcal/mol and 0.8 to 1.3 kcal/mol M^−1^, respectively (Table 1)^35–37^. Notably, both the apparent stability and *m*-value of WholeCons* Core are similar to the calculated apparent stability and *m*-value of the entire WholeCons*. Together, these data suggest that the consensus approach may have resulted in a biased stabilization of the core region of RNase H, such that the periphery contributes little to the overall stability, modifying the typical equilibrium two-state behavior of RNases H.

### Inclusion of additional C-terminal residues does not recover two-state behavior to WholeCons*

Extant RNase H sequences vary in their length, mostly due to the variable inclusion of several C-terminal residues, called ‘tails’. *E. coli* RNase H has such a tail (residues 145 – 154) and these C-terminal residues are known to affect the stability of ecRNH^38, 39^. WholeCons* lacks the ten C-terminal residues present in ecRNH. We evaluated several additional constructs and confirmed the lack of these C-terminal residues is not responsible for the apparent non-cooperative behavior (see Supplement 1).

### The thermodynamic properties of RNase H consensus proteins are sensitive to the diversity of the input sequences

To examine how the properties of consensus proteins depend on the number and diversity of the input sequences, we generated several additional consensus proteins (Anc1cons, AncAcons, AncCcons*, DispersedCons*). As discussed above, these sequences vary in the evolutionary similarity and number of input sequences. All express and purify similarly to other RNases H, are enzymatically active, and have CD spectra typical of folded RNase H (for example, see Figure 2). The stabilities of these variants were investigated using the same chemical and thermal denaturation and two-state assumption discussed above (Figure 3C,D, Table 1). Consensus variants derived from sequences spanning the RNase H phylogenetic tree (Anc1cons, AncAcons, and DispersedCons*), behave similarly to WholeCons* showing low apparent stabilities and cooperativities (Table 1).

In contrast, AncCcons*, which is derived from closely related sequences, behaves more like a typical RNase H with an *m*-value consistent with that expected for the RNase H fold (2.1 ± 0.1 kcal/mol M^−1^). In addition, AncCcons* shows a higher apparent stability compared to the other consensus proteins: ΔG_unf_(25 °C) = 11.8 ± 0.6 kcal/mol, higher than most of the ancestors and close to that of the thermophilic ttRNH. The T_m_ of AncCcons* is 76.3 ± 2.3 °C, 7.9 to 24.6 °C greater than RNases H in the clade it is derived from (Table 2)^23^. Figure 6 summarizes the resulting T_m_s of all the consensus proteins evaluated.

**Figure 6.**
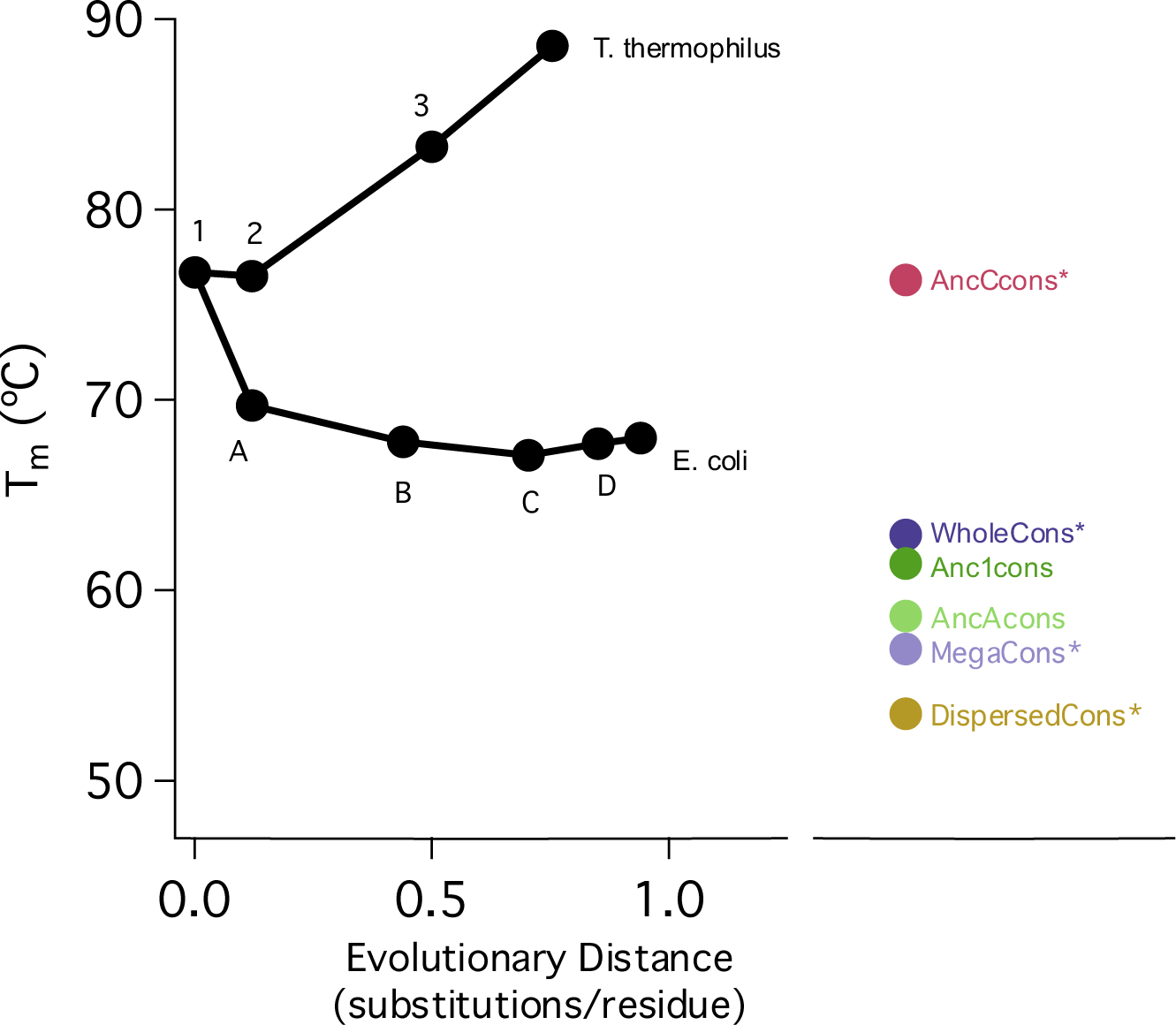
Summary of melting temperatures for consensus proteins compared to ancestral and extant RNases H^23^.

**Table 2:** T_m_s of AncC descendants^23^ compared to AncCcons*.

| | T<sub>m</sub> (°C) | $\Delta T_m$ from AncCcons* (°C) | %ID to AncCcons* |
| --- | --- | --- | --- |
| *Escherichia coli* | 68.0 ±0.5 | -8.3 | 88.96 |
| *Klebsiella pneumoniae* | 68.4 ±0.3 | -7.9 | 89.61 |
| *Citrobacter sp 30_2* | 62.4 ±0.4 | -13.9 | 85.71 |
| *Cronobacter turicensis* | 63.8 | -12.5 | 88.89 |
| *Salmonella enterica* | 59.1 | -17.2 | 88.36 |
| *Enterobacter sp. 638* | 63.1 ±0.1 | -13.2 | 85.06 |
| *C. hamiltonella defensa* | 51.7 ±1.1 | -24.6 | 75.82 |

Errors reported are standard deviations from replicate experiments.

The increased *m*-value of AncCcons* suggests the protein behaves more like a typical extant RNase H, with a cooperative two-state folding equilibrium folding mechanism and a kinetic folding trajectory that shows a burst-phase intermediate. To probe this further, we studied the folding kinetics of AncCcons*. The unfolding and refolding kinetics showed an early burst-phase intermediate followed by a single exponential phase (Figure 7). The resulting chevron plot shows a roll-over at low denaturant concentrations, indicating that AncCcons* folds by a similar three-state kinetic folding mechanism observed for the extant RNases H (Figure 7 and Table 3). In sum, these data suggest that AncCcons*, which is derived from sequences that are in a restricted part of the phylogenetic tree, is a cooperatively folded and stabilized consensus RNase H.

**Figure 7.**
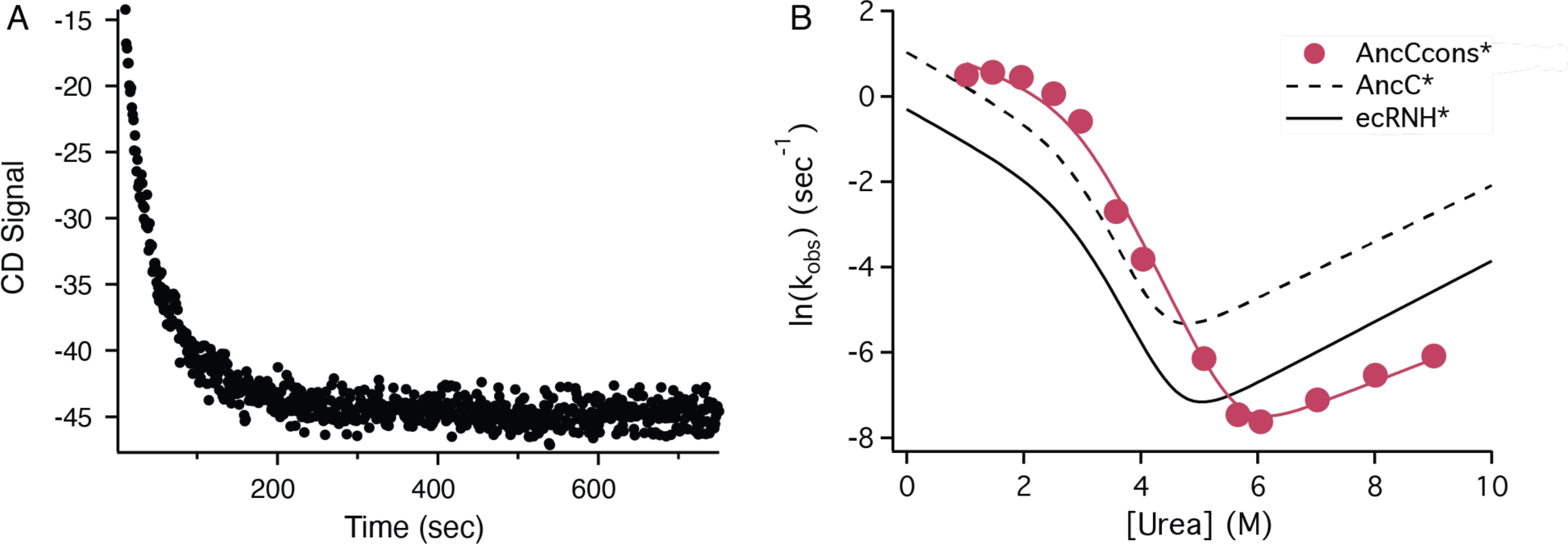
Kinetics of AncCcons* folding. **(A).** Refolding of AncCcons* at 4M Urea monitored by CD at 222 nm, showing single exponential decay. **(B).** Chevron plot of AncCcons* compared to fitted cures for AncC* and ecRNH*^24^

**Table 3:** Kinetic (three-state kinetic folding model) parameters for AncCcons*, AncC*^24^, and ecRNH*^18^.

| | $k_{in}$<br>(s<sup>-1</sup>) | $m_{in}$<br>(kcal/mol/M) | $k_{ni}$<br>(s<sup>-1</sup>) | $m_{ni}$<br>(kcal/mol/M) | $\Delta G_{UI}$<br>(kcal/mol) | $m_{UI}$<br>(kcal/mol/M) | $\Delta G_{unf}$<br>(kcal/mol) | $m_{unf}$<br>(kcal/mol/M) |
| --- | --- | --- | --- | --- | --- | --- | --- | --- |
| AncCcons* | 3.8<br>±2.4 | 0.31 ±0.17 | 1.8 ±1.9 × 10<sup>-5</sup> | -0.31 ±0.09 | 4.0 ±4.5 | 1.35 ±0.21 | 11.3 ±4.5 | 1.97 ±0.28 |
| AncC* | 2.8<br>±0.9 | 0.47 ±0.10 | 1.7 ±0.6 × 10<sup>-4</sup> | -0.39 ±0.04 | 3.9 ±0.9 | 1.34 ±0.17 | 9.7 ±1.0 | 2.23 ±0.20 |
| ecRNH | 0.74<br>±0.02 | 0.45 ±0.01 | 1.7 ±0.1 × 10<sup>-5</sup> | -0.42 ±0.03 | 3.5 ±0.1 | 1.24 ±0.08 | 9.9 ±0.1 | 2.11 ±0.09 |

### Evolutionary couplings are equally maintained for all RNase H consensus proteins

Recent computational analyses of MSAs highlight the importance of evolutionary couplings between specific sites in a protein family^40, 41^. These evolutionary couplings are observed as a statistical propensity for two positions to co-evolve, particularly for positions that are adjacent in the three-dimensional structure, and are measured as a covariance. Given an MSA, one can calculate covariance by comparing observed pair frequencies (e.g., the frequency of both residue A at position i and residue B occurs in position j) with the product of the frequencies of residue A in Position 1 and residue B in Position 2). If there is positive covariance, the observed pair frequency will be higher than the product of the singe-residue frequencies. We set out to understand how evolutionary coupling is captured (or not captured) by the consensus and ancestral RNases H.

Because consensus proteins are generated by simply counting the frequency at each site independent of any other site, favorable coupling between pairs of residues in extant sequences may be lost in the resulting consensus sequences, though they are expected to be retained in ancestral sequences. To investigate whether the observed loss in cooperativity and stability in some of our consensus sequences results from the omission of favorable pairwise coupling, we quantified the pairwise couplings in the RNase H family using a Potts model^42, 43^. In this model, the intrinsic energy of each possible amino acid at each site (*H_i_*) and the coupling energy between each possible pair of amino acids at each pair of sites (*J_ij_*) can be estimated from a large MSA using maximum likelihood optimization. To help capture weakly coupled residue pairs, we expanded our number of RNase H sequences from 405 to ∼11,300 using the InterPro database (see Methods, Figure 8). With this larger MSA, similar *H_seq_* and *J_seq_* values are obtained when the 11,300 sequence MSA is randomly split in half, indicating that the sequence set is large enough to reliably estimate *J_seq_* values.

**Figure 8.**
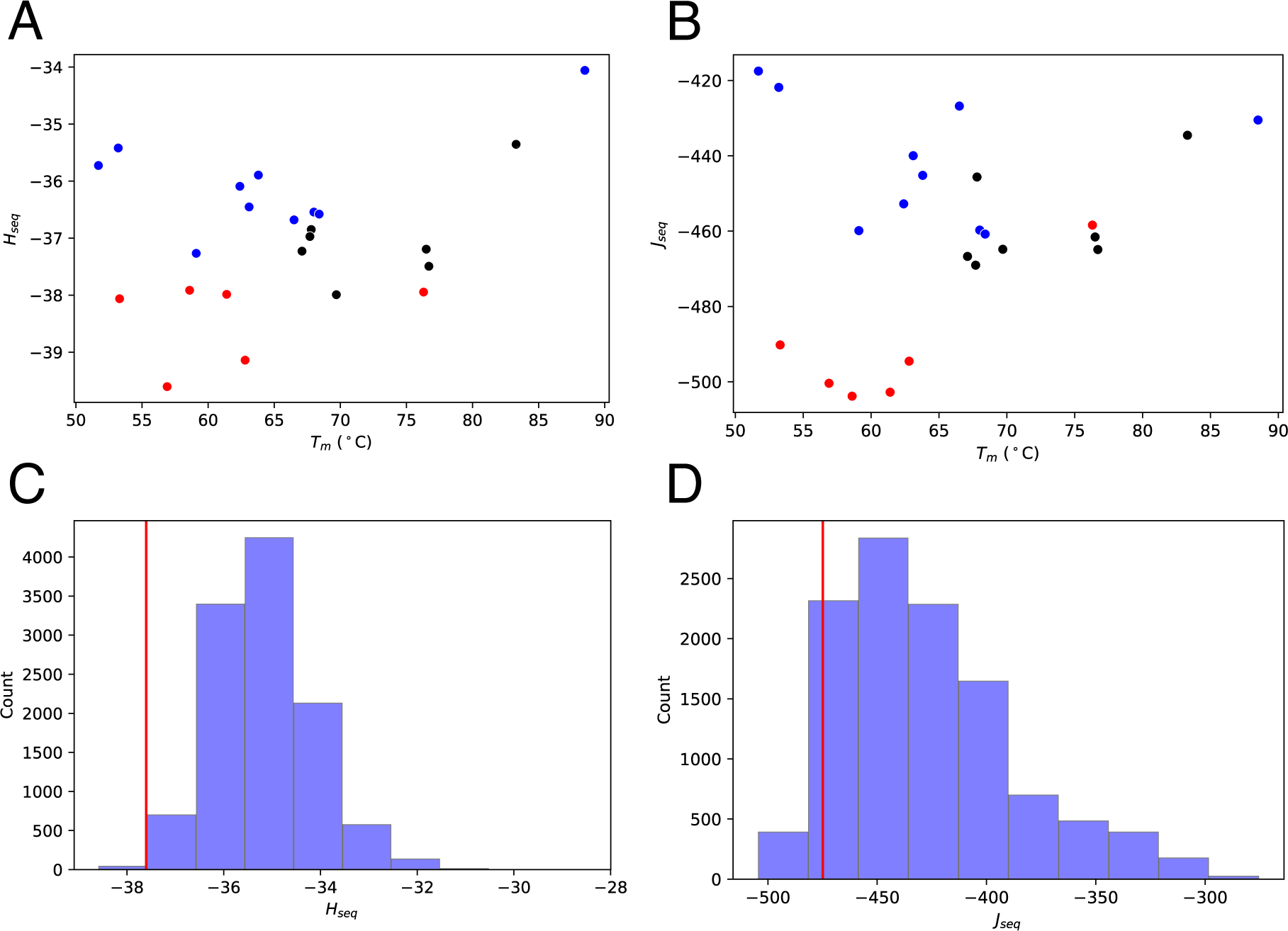
Potts analysis of RNaseH sequences. Intrinsic and pairwise coupling coefficients were determined from an MSA with 11,300 sequences, and were used to calculate total intrinsic (*H_seq_*, **A**) and pairwise coupling scores (*J_seq_*, **B**) for consensus (red), ancestral (black), and extant (blue) sequences. Neither score correlates strongly with overall stability, as represented with T_m_ values. **(C, D)** Histograms of *H_seq_* and *J_seq_* scores for the 11,300 extant sequences in the alignment, along with the values from the consensus derived from those 11,300 sequences (red lines).

Once we had reliable estimates for *H_i_* and *J_ij_*, we used them to calculate the total intrinsic energy (*H_seq_*) and total coupling energy (*J_seq_*) for each of the ancestral, consensus, and extant sequences. These can be calculated for a sequence by a simple sum over all residues and pairs:

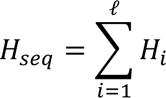

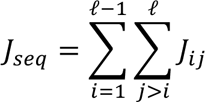

where ℓ is the length of the MSA and *H_i_* and *J_ij_* are the coefficients estimated from the large MSA. Here we take favorable Potts *H* and *J* terms (i.e., strong conservation and positive covariance respectively) to be negative in sign to provide an energetic perspective.

The ancestral and consensus sequences have a range of *H_seq_* and *J_seq_* values (Figure 8). Not surprisingly, the consensus sequences (red circles, Figure 8A) have the lowest (i.e., most stabilizing) average *H_seq_* values, which is consistent with the fact that the *H_i_* coefficients are related to the extent of site-specific conservation. In comparison, extant sequences for which T_m_ values are known (blue circles in Figure 8A) have the highest average *H_seq_* values. These are consistent with the overall distribution of *H_seq_* values for the 11,300 interpro sequences, which have considerably higher *H_seq_* values than that of the interpro consensus (red line, Figure 8C). The ancestral sequences have *H_seq_* scores that are, on average, in between those of the consensus and extant sequences, although there is some overlap between the ancestral and extant sequences (Figure 8A), suggesting that ancestral sequences are more similar to extant sequences than are consensus sequences.

Similarly, though perhaps unexpectedly, with one exception the consensus sequences also have the lowest *J_seq_* values (Figure 8B). Again, the ancestral sequences have *J_seq_* values that are in between those of the consensus and extant sequences. This pattern is inconsistent with the hypothesis that the observed loss in cooperativity and stability in some of our consensus sequences results from the omission of favorable pairwise couplings. To underscore this observation, the AncCcons* sequences has the highest T_m_ and cooperativity of the consensus sequences studied here, and it has the highest (least stabilizing in terms of overall Potts energy) *J_seq_* value. Moreover, extant RNaseH sequences that span the entire range of observed T_m_ values have identical *J_seq_* values.

### Analysis of ancestral and derived consensus sequences in SVD space

The studies here demonstrate that for the RNase H family, the properties of consensus designs from sequences that descend from ancestral nodes in the RNase H tree differ from the ancestral sequences themselves. To better understand these stability differences, and why the consensus sequences derived from the deep Anc1 and AncA nodes are destabilized, whereas the consensus derived from AncC is stabilized, we examined the relative positions of these sequences in singular value decomposition space.

Singular value decomposition (SVD) is an approach for “dimensionality reduction” in the same vein as the more familiar Principal Component Analysis. SVD reduces each sequence to a point in a high-dimensional space. The axes of this space are organized from the highest to lowest contribution to the observed variance. One can compare sequences by asking the extent to which they spread out and/or cluster with one another along the first few axes of the SVD.

Mathematically, SVD takes a rectangular matrix *F* and transforms it to a product of two orthogonal matrices *U* and *V* that are related to the rows and columns of the original matrix. SVD can be applied to a multiple sequence alignment, which can be viewed as a matrix where the rows are sequences and the columns are residues^44, 45^. By first encoding each position in the MSA using a binary encoding, SVD produces a coordinate system in which each sequence in the MSA can be depicted as a single point on axes denoted *σ_1_u_i_^(1)^*, *σ_2_u_i_^(2)^*, … *σ_N_u_i_^(N)^*, where *N* is the number of sequences in the MSA (409 for the MSA used in ancestral reconstruction). One of the properties of SVD is that the most important variances and covariances are concentrated into a small number of coordinates, allowing clusters of sequences to be resolved and their differences to be visualized in sequence space in a quantitative way. Moreover, by alignment to the MSA, sequences not in the original MSA (such as ancestors and consensus sequences) can be projected into SVD space^46^.

To visualize the relationships among the extant, ancestral, and consensus RNase H sequences, we performed SVD on the same MSA used for ancestral reconstruction (the 405 sequences in the tree plus the four used to root the tree) and projected the ancestral and consensus sequences into this space along with the sequences used for the SVD (Figure 9). As we have found with other sequence families, the RNase H MSA sequences cluster into distinct groups. These groups can be highlighted using a *k*-means algorithm (Figure 9A) with five sequence clusters. Sequences that descend from Anc1 and AncA (which are nearly the same sequences — the Anc1 descendants have 5 sequences in addition to the 298 sequences in AncA) are composed of two distinct clusters with very different *σ_2_u_i_^(2)^* values (Figure 9B and C). The first, which includes the yellow and green *k*-means clusters (173 and 53 sequences respectively), corresponds to descendants from AncA (and the similar Anc1 group) that do not include AncB and its descendants (gold, Figure 9E). The second Anc1/AncA cluster, which includes the blue *k*-means cluster (79 sequences), corresponds to descendants from AncB. The bottom tip of the blue cluster, at an extreme negative value of *σ_3_u_i_^(3)^* corresponds to descendants of the AncC cluster (Figure 9D). The overlapping patterns of SVD clusters and ancestral descendants underscores the close connection between sequence phylogeny and patterns in SVD space.

**Figure 9.**
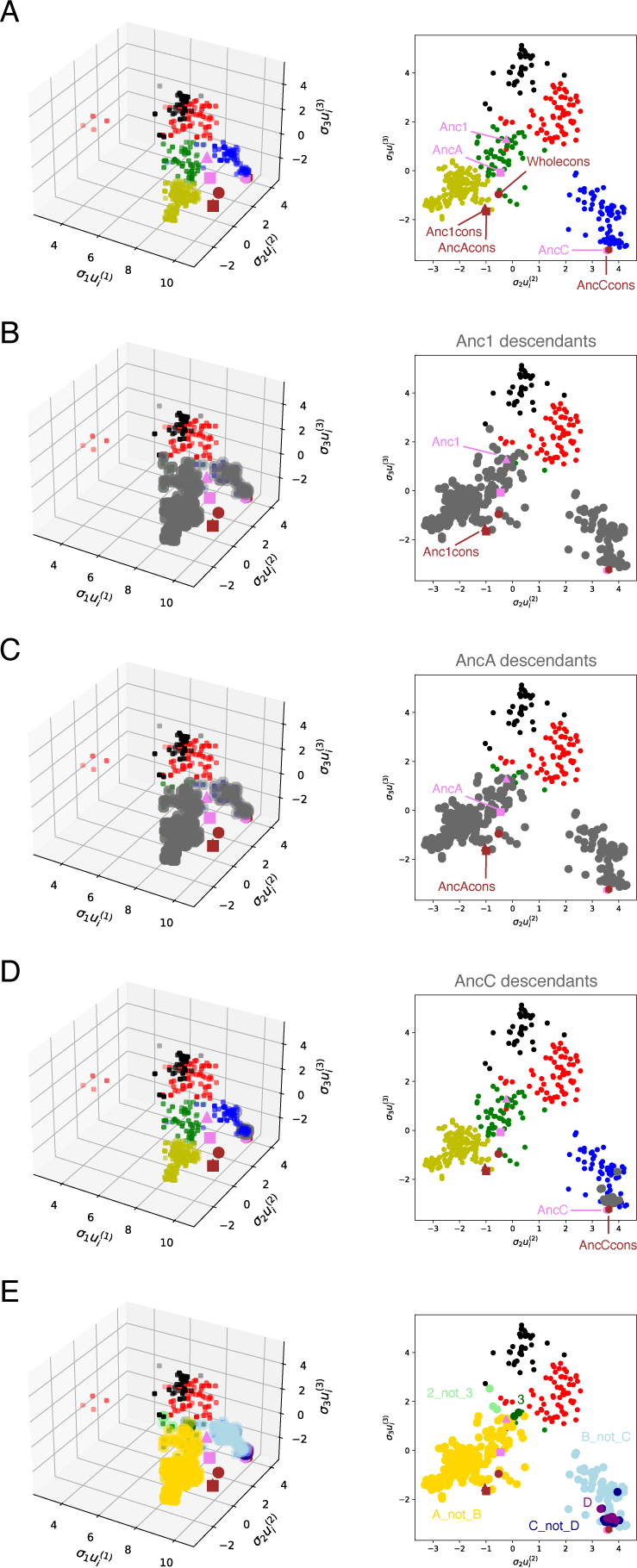
RNase H sequences in SVD space. 409 aligned RNase H sequences from Hart et al^23^ were represented as a binary matrix, which was transformed into an orthogonal sequence and residue matrices using singular value decomposition^46^. Plots show sequences (small points) plotted in the first three dimensions of SVD space (left), and in dimensions 2 and 3 (right). The first dimension (*σ_1_u_i_^(1)^*), shown only in the three-dimensional plots, reflects overall conservation, whereas the second and third dimensions reflect covariance patterns among sequences. (**A**) MSA sequences used in the SVD are colored red, black, blue, green, and yellow based on *k*-means clustering into five groups. Ancestral sequences projected into SVD space are colored violet, and consensus sequences created from ancestral descendant sequences are colored brown. (**B-D**) Descendants of the main ancestors investigated here (Anc1, AncA, and AncC, respectively) are plotted as large grey spheres. Descendants of AncC are also descendants of AncA (and Anc1), and those of AncA are also descendants of Anc1, resulting in considerable overlap. (**E**) Non-overlapping descendants of major ancestral branchpoints, plotted as large colored spheres. The python scripts that generated the binary encoding, SVD, and plots are available as a jupyter notebook at https://github.com/barricklab-at-jhu/SVD-of-MSAs/tree/main/RNaseH“.

Projecting the ancestral and derived consensus sequences into SVD space reveals interesting similarities and differences among these sequences, as well as relationships of ancestors and consensus sequences to SVD clusters. When viewed projected onto the 2^nd^ and 3^rd^ axes (the *σ_2_u_i_^(2)^ σ_3_u_i_^(3)^* plane), sequences from the phylogenetically restricted “C” group cluster together. This includes the AncC sequence, the AncCcons, and the extant sequences that descend from AncC (Figure 9A, D). In contrast, the consensus sequences derived from the larger Anc1 and AncA descendants are quite distant from the descendants themselves (Figure 9A-C). Specifically, Anc1cons and AncAcons are located between the yellow, green, and blue *k*-means clusters (all three are of which are descendant from Anc1/AncA), lying nearest to the yellow group (descendants of AncA that do not include those of AncB). Proximity of the Anc1/A consensus to the yellow cluster is to be expected, since the yellow cluster has more sequences than the green or blue clusters, and thus should have more impact on the derived consensus sequences. In contrast, the ancestral sequences AncA and Anc1 do not group with the derived consensus sequences, but instead are positioned near the green cluster in the *σ_2_u_i_^(2)^ σ_3_u_i_^(3)^* plane.

The locations in SVD space of the ancestral and derived consensus sequences relative to the extant RNase H sequences provides a rationalization for the stability patterns described above: consensus sequences achieve high stability and cooperativity when they retain positions in SVD space that reflect the sequence covariance of the extant sequences (e.g., they group with the extant sequences in *σ_2_u_i_^(2)^*, *σ_3_u_i_^(3)^*, and perhaps higher dimensions). In other words, successful consensus design requires some threshold level of similarity to the correlations seen in extant sequences in the main SVD coordinates that capture correlation. Importantly, these coordinates, which capture large covariations among multiple positions, have been shown to discriminate among subfamilies of sequences with deep phylogenetic divergence^47^. Consensus sequences generated from widely different subfamilies, such as Anc1cons and AncAcons, can be quite different from all of the extant sequences in terms of their covariance patterns, and may have low stability as a result, even though their residue compositions match those in the MSA. This stability pattern is reminiscent of that seen by Magliery and coworkers, who found that consensus TIM barrels from taxonomically restricted MSAs were better behaved and more stable than those from a broad MSA^13^.

AncCcons, which is highly stabilized and folds cooperatively, is close to the AncC descendants from which it was designed in the *σ_2_u_i_^(2)^ σ_3_u_i_^(3)^* plane. In contrast, Anc1cons and AncAcons are distant from most of the sequences from which they were designed, and have decreased stability. The ancestral sequences for these two groups, which are stable and cooperative, are far from their consensus sequences in the *σ_2_u_i_^(2)^ σ_3_u_i_^(3)^* plane, and are located more centrally within the green cluster. It is noteworthy that the coordinates of Anc1cons and AncAcons in the first dimension (*σ_1_u_i_^(1)^*, a measure of overall conservation) are larger than the coordinates of Anc1 and AncA, yet they are less stable, likely a reflection of their positioning in the *σ_2_u_i_^(2)^ σ_3_u_i_^(3)^* plane (and in perhaps in higher dimension).

Finally, the differences in *σ_2_u_i_^(2)^ σ_3_u_i_^(3)^* coordinates between the Anc1 and AncA sequences and their corresponding consensus sequences illustrates a fundamental difference between ASR and consensus design. By nature, consensus design generates sequences with “average” properties, and this averaging extends into the covariances that are captured in higher-order SVD coordinates. This averaging can produce sequences that have significant differences from all natural sequences, extant or extinct, and as a result, may be compromised in stability or function. Moreover, the Potts analysis (Figure 8) shows that consensus sequences can have large negative (i.e., stabilizing in terms of overall Potts energy) pairwise covariance scores, even though they miss the mark in terms of the higher order correlations represented by SVD. In contrast, ancestral sequences were shaped by natural selection, and will occupy a region of sequence space that encodes stability and function. For RNase H, that region appears to have been the region occupied by extant sequences in the green k-means cluster, which generated green descendants locally, as well as more distant yellow and blue descendants.

## Discussion

Here, we show that ancestral and consensus proteins can have very different biophysical properties. For RNase H, ancestral proteins Anc1 and AncA have relatively high T_m_ values of 76.7 and 69.7 °C, whereas the consensus proteins Anc1cons and AncAcons have low T_m_ values of 61.4 and 58.6 °C, and show broad urea-induced unfolding transitions. In contrast, the consensus protein derived from the AncC (AncCcons) is significantly more stable, with a T_m_ of 76.3 °C (9.2 °C higher than AncC), and has a sharp urea-induced unfolding transition, demonstrating that the properties of consensus proteins are sensitive to the phylogenetic relationship of input sequences.

Since the consensus method is a simple counting method at each position, the resulting consensus sequence is likely to be sensitive to the number and diversity of the input sequences. This is in contrast to ASR, where it has been well-established that the inferred ancestral sequences are fairly robust to the diversity of input sequences^48^. ASR considers the entire topology and content of the tree to build an ancestor at a given node. Here, we demonstrate the difference in the two methods by directly comparing the consensus and ancestral proteins derived from the same clades. The set of 303 and 24 extant sequences descending from Anc1 and AncC, respectively, were used to generate a consensus protein. If the two methods - ASR and consensus - generate similar proteins, we would expect the properties of Anc1 to be similar to Anc1cons and the properties of AncC to be similar to AncCcons*. However, we do not find this to be the case. The T_m_s of Anc1 and Anc1cons differ by 15.3 ± 1 °C, with the ancestor being more stable. Conversely, AncC and AncCcons* differ by 9.2 ± 2.3 °C, with the consensus protein being more stable. No consistent trend is found when comparing these ancestral and consensus RNase H proteins, further distinguishing the two methods.

Importantly, whereas the ancestral sequences produce cooperatively folded proteins, such as Anc1*, the consensus method usually results in non-cooperative multi-state folding, such as WholeCons*. Several of the 32 mutations between ecRNH and WholeCons* reside at key locations in the interface where the periphery of the protein docks onto the core, which takes place during the rate-limiting step from the kinetic intermediate to the native state. These mutations may increase the folding barrier between the core intermediate and native state or decrease the stability of the native state.

The population of a partially unfolded equilibrium intermediate under native conditions as observed in WholeCons* is expected to affect the binding strength and specificity for different binding partners as well as enzymatic activity. Such partially unfolded intermediates will be more susceptible to proteolytic degradation and aggregation than the native protein. The higher *m*-value seen in the consensus protein derived from an evolutionarily related set (AncCcons) suggests that evolution drives RNase H proteins to maintain cooperative folding behavior, possibly by a variety of mechanisms that are disrupted when evolutionarily diverse sequences are used to design a consensus. Cooperativity may be encoded by different mechanisms in separate clades of the tree and is thus lost when too many diverse clades are combined to generate a consensus protein.

This study highlights a case where the consensus method does not always increase protein stability and can disrupt cooperative folding. Most previously published consensus work supports the idea that the consensus method is a generalizable approach to create thermostable proteins. While RNase H may be an outlier, it is possible that our results are less exceptional than they seem, since unstable and poorly behaved consensus-designed proteins are unlikely to be published.

Our SVD analysis points to potential fundamental differences in ASR and consensus design. Along with the differences in stability, we found differences in *σ_2_u_i_^(2)^ σ_3_u_i_^(3)^* coordinates between the Anc1/AncA and their corresponding consensus sequences. By nature, consensus design generates sequences with “average” properties, and this averaging extends into the covariances that are captured in higher-order SVD coordinates. This averaging can produce sequences that have significant differences from all natural sequences, extant or extinct, and as a result, may be compromised in stability or function. In contrast, ancestral sequences were shaped by natural selection, and will occupy a region of sequence space that encodes stability and function. If the results shown here are generalizable, it suggests an approach where SVD can be used to refine consensus design by restricting it to regions of SVD space populated by extant (and perhaps ancestral) sequences.

## Methods

### Consensus Design

WholeCons*, Anc1cons, AncAcons, and AncCcons* protein sequences were designed by taking the most common amino acid at each position of the MSA of the RNase H family^23^. MegaCons* was generated from sequences obtained from InterPro^49^ using Ribonuclease HI family (accession number IPR22892) as a query, with domain architecture that only included a single RNase HI domain, culling sequences with greater than 0.7x or less than 1.2x of 155 residues (median length from set) and CD-HIT 0.9 threshold sequence identity^50^. The resulting 3,312 sequences were aligned with MAFFT^51^ and MSA_analysis.py script was used to generate a residue frequency matrix. The most frequent amino acid at each position was used to generate the final MegaCons* sequence.

### Consensus Expression and Purification

Integrated DNA Technologies (IDT) gBlock consensus RNase H gene fragments were restriction cloned into the pET-27b(+) vector for expression. Site-directed mutagenesis was used to produce consensus C-terminal tail variants. The sequences were verified by sanger sequencing. The proteins were expressed and purified as described^17^. The purity and mass of the proteins were confirmed by SDS/PAGE and mass spectrometry.

### Circular Dichroism Spectroscopy

CD experiments were measured using an Aviv 410 CD spectrometer. For spectra, signals from 200 nm to 300 nm were obtained for samples of 0.4 mg/mL protein in 20 mM sodium acetate and 50 mM potassium chloride, pH 5.5 (RNase H buffer) using a 1-mm path length Starna Cells cuvette at 25 °C. The samples were buffer corrected by subtracting the signal of buffer without protein at each wavelength. For urea melts, samples containing 40 μg/mL protein and varying [urea] in RNase H buffer were equilibrated and the signal at 222 nm was measured in a 1-cm path length Starna Cells cuvette at 25 °C with stirring, averaged over one minute. For temperature melts, data was collected at 222 nm every 3 °C on samples containing 40 µg/mL protein with a path length of 1-cm, averaged over one minute. Samples were equilibrated while stirring for 5 min at each temperature. Anc1cons and AncAcons experiments were performed in RNase H buffer with the addition of 1 mM TCEP. Melts were fit to a two-state model with linear free-energy extrapolation and the apparent T_m_ was calculated in IgorPro.^25^ CD melts were obtained in triplicate.

### RNase H Activity Assay

RNase H activity was measured with a substrate prepared from equal parts dT20 oligomers (IDT) and poly-rA (Sigma), heated to 95 °C for 5 minutes, followed by slow cooling to room temperature for 1 hour. The buffer conditions were 10 mM Tris, 50 mM NaCl, 10 mM MgCl_2_, 1 mM TCEP, pH 8, at 25 °C. The reaction was initiated by the addition of 65 nM enzyme and monitored at 260 nm using a Cary UV spectrophotometer.

### Fluorescence Spectroscopy

Fluorescence measurements were obtained on a FluoroMax-3 fluorometer. For spectra, the same urea melt samples from CD experiments were excited at 295 nm and emission was acquired every 1nm from 305 nm to 400 nm. The ratio of 374/324 nm was plotted vs. urea and the resulting curve was fit as described for CD melts. Fluorescence melts were obtained in duplicate.

### Kinetic analysis

RNase H folding kinetics were performed on an Aviv stopped-flow circular dichroism spectrometer, model 202, as previously described on ancestral RNases H.^24^

### Pulse-labeling hydrogen-deuterium exchange monitored by mass spectrometry (HXMS)

HXMS experiments were performed as previously described.^33^

### DCA Potts model analysis

Using the InterPro database, all unique bacterial RNases H were collected and pruned to only include sequences that are in isolated domains, a length +/− 30% of median: 154 amino acids. All positions with less than 80% gap frequency were kept in the resulting MSA. The DCA Potts model was fit as described^42^ using the default settings of the code.

### Single Value Decomposition

SVD was carried out as detailed previously^46^.

## Acknowledgements

We thank the entire Marqusee and Barrick labs and Katie Tripp for help and advice. Special thanks to Mark Petersen for critical reading of the manuscript. The work was funded by grants from the NIH (GM050945, SM) and (GM068462, DB). SM is a Chan Zuckerberg Biohub Investigator.

## References

1. Magliery TJ (2015) Protein stability: Computation, sequence statistics, and new experimental methods. Curr. Opin. Struct. Biol. 33:161–168.

2. Porebski BT, Buckle AM (2016) Consensus protein design. Protein Eng. Des. Sel. 29:245– 251.

3. Thornton JW, Need E, Crews D (2003) Resurrecting the ancestral steroid receptor: Ancient origin of estrogen signaling. Science (80-.). 301:1714–1717.

4. Wilson C, Agafonov R V., Hoemberger M, Kutter S, Zorba A, Halpin J, Buosi V, Otten R, Waterman D, Theobald DL, et al. (2015) Using ancient protein kinases to unravel a modern cancer drug’s mechanism. Science (80-.). 347:882–886.

5. Anderson JA, Loes AN, Waddell GL, Harms MJ (2019) Tracing the evolution of novel features of human Toll-like receptor 4. Protein Sci. 28:1350–1358.

6. Merkl R, Sterner R (2016) Ancestral protein reconstruction: Techniques and applications. Biol. Chem. 397:1–21.

7. Hochberg GKA, Thornton JW (2017) Reconstructing Ancient Proteins to Understand the Causes of Structure and Function. Annu. Rev. Biophys. 46:247–269.

8. Di Giulio M (2003) The universal ancestor was a thermophile or a hyperthermophile: Tests and further evidence. J. Theor. Biol. 221:425–436.

9. Akanuma S, Nakajima Y, Yokobori S, Kimura M, Nemoto N, Mase T (2013) Experimental evidence for the thermophilicity of ancestral life. PNAS 110:1–6.

10. Sternke M, Tripp KW, Barrick D (2019) Consensus sequence design as a general strategy to create hyperstable, biologically active proteins. Proc. Natl. Acad. Sci. U. S. A. 166:11275– 11284.

11. Risso VA, Sanchez-Ruiz JM, Ozkan SB (2018) Biotechnological and protein-engineering implications of ancestral protein resurrection. Curr. Opin. Struct. Biol. 51:106–115.

12. Sullivan BJ, Nguyen T, Durani V, Mathur D, Rojas S, Thomas M, Syu T, Magliery TJ (2012) Stabilizing proteins from sequence statistics: The interplay of conservation and correlation in triosephosphate isomerase stability. J. Mol. Biol. 420:384–399.

13. Goyal, VD; Sullivan, BJ; Magliery T (2019) Phylogenetic Spread of Sequence Data Affects Fitness of Consensus Enzymes: Insights from Triosephosphate Isomerase. Proteins Struct. Funct. Bioinforma.

14. Nakano S, Motoyama T, Miyashita Y, Ishizuka Y, Matsuo N, Tokiwa H, Shinoda S, Asano Y, Ito S (2018) Benchmark Analysis of Native and Artificial NAD+-Dependent Enzymes Generated by a Sequence-Based Design Method with or without Phylogenetic Data. Biochemistry 57:3722–3732.

15. Tadokoro T, Kanaya S (2009) Ribonuclease H: Molecular diversities, substrate binding domains, and catalytic mechanism of the prokaryotic enzymes. FEBS J. 276:1482–1493.

16. Dabora JM, Marqusee S (1994) Equilibrium unfolding of Escherichia coli ribonuclease H: Characterization of a partially folded state. Protein Sci. 3:1401–1408.

17. Raschke TM, Marqusee S (1997) The kinetic folding intermediate of ribonuclease H resembles the acid molten globule and partially unfolded molecules detected under native conditions. Nat. Struct. Biol. 4.

18. Raschke TM, Kho J, Marqusee S (1999) Confirmation of the hierarchical folding of RNase H: a protein engineering study. Nat Struct Biol 6:825–831.

19. Hollien J, Marqusee S (1999) A thermodynamic comparison of mesophilic and thermophilic ribonucleases H. Biochemistry 38:3831–3836.

20. Hollien J, Marqusee S (1999) Structural distribution of stability in a thermophilic enzyme. Proc. Natl. Acad. Sci. U. S. A. 96:13674–13678.

21. Hollien J, Marqusee S (2002) Comparison of the folding processes of T. thermophilus and E. coli ribonucleases H. J. Mol. Biol. 316:327–40.

22. Robic S, Berger JM, Marqusee S (2002) Contributions of folding cores to the thermostabilities of two ribonucleases H. Protein Sci. 11:381–9.

23. Hart KM, Harms MJ, Schmidt BH, Elya C, Thornton JW, Marqusee S (2014) Thermodynamic system drift in protein evolution Theobald DL, editor. PLoS Biol 12:e1001994.

24. Lim SA, Hart KM, Harms MJ, Marqusee S (2016) Evolutionary trend toward kinetic stability in the folding trajectory of RNases H. Proc. Natl. Acad. Sci. U. S. A. 113:13045–13050.

25. Street TO, Courtemanche N, Barrick D (2008) Protein Folding and Stability Using Denaturants. Methods Cell Biol. 84:295–325.

26. Tanford C (1970) Protein denaturation: Part c. theoretical models for the mechanism of denaturation. Adv. Protein Chem. 24:1–95.

27. Shortle D (1995) Staphylococcal Nuclease: A Showcase of m-value effects. Adv. Protein Chem. 46.

28. Myers JK, Nick Pace C, Martin Scholtz J (1995) Denaturant m values and heat capacity changes: Relation to changes in accessible surface areas of protein unfolding. Protein Sci. 4:2138–2148.

29. Creighton TE, Shortle D (1994) Electrophoretic characterization of the denatured states of staphylococcal nuclease. J. Mol. Biol. 242:670–682.

30. Soulages JL (1998) Chemical denaturation: Potential impact of undetected intermediates in the free energy of unfolding and m-values obtained from a two-state assumption. Biophys. J. 75:484–492.

31. Spudich G, Marqusee S (2000) A change in the apparent m value reveals a populated intermediate under equilibrium conditions in Escherichia coli ribonuclease HI. Biochemistry 39,38 11677–11683

32. Hu W, Walters BT, Kan Z-YY, Mayne L, Rosen LE, Marqusee S, Englander SW (2013) Stepwise protein folding at near amino acid resolution by hydrogen exchange and mass spectrometry. Proc Natl Acad Sci U S A 110:7684–7689.

33. Lim SA, Bolin ER, Marqusee S (2018) Tracing a protein’s folding pathway over evolutionary time using ancestral sequence reconstruction and hydrogen exchange. Elife 7:38369

34. Chamberlain AK, Fischer KF, Reardon D, Handel TM, Marqusee S (1999) Folding of an isolated ribonuclease H core fragment. Protein Sci. 8:2251–2257.

35. Rosen LE, Connell KB, Marqusee S (2014) Evidence for close side-chain packing in an early protein folding intermediate previously assumed to be a molten globule. Proc. Natl. Acad. Sci. U. S. A. 111:14746–51.

36. Rosen LE, Marqusee S (2015) Autonomously folding protein fragments reveal differences in the energy landscapes of homologous rnases He0119640. PLoS One 10:1–14.

37. Lim SA, Marqusee S (2018) The burst-phase folding intermediate of ribonuclease H changes conformation over evolutionary history. Biopolymers 109:e23086.

38. Haruki M, Noguchi E, Akasako A, Oobatake M, Itaya M, Kanaya S (1994) A novel strategy for stabilization of Escherichia coli ribonuclease HI involving a screen for an intragenic suppressor of carboxyl-terminal deletions. J. Biol. Chem. 269:26904–26911.

39. Goedken ER, Raschke TM, Marqusee S (1997) Importance of the C-terminal helix to the stability and enzymatic activity of Escherichia coli ribonuclease H. Biochemistry 36:7256– 7263.

40. Hopf TA, Ingraham JB, Poelwijk FJ, Springer M, Sander C, Marks DS (2015) Quantification of the effect of mutations using a global probability model of natural sequence variation. arXiv preprint arXiv:151004612.

41. Hopf TA, Green AG, Schubert B, Mersmann S, Schärfe CPI, Ingraham JB, Toth-Petroczy A, Brock K, Riesselman AJ, Palmedo P, et al. (2019) The EVcouplings Python framework for coevolutionary sequence analysis. Bioinformatics 35:1582–1584.

42. Ekeberg M, Cecilia L, Lan Y, Weigt M, Aurell E (2013) Improved contact prediction in proteins: Using pseudolikelihoods to infer Potts models. Phys. Rev. E 87, 012707:1–16.

43. Tian P, Louis JM, Baber JL, Aniana A, Best RB (2018) Co-Evolutionary Fitness Landscapes for Sequence Design. Angew. Chemie - Int. Ed.:5674–5678.

44. Casari G, Sander C, Valencia A (1995) A method to predict functional residues in proteins. Nat. Struct. Biol. 1995 22 2:171–178.

45. Gogos A, Jantz D, Sentü S, Richardson D, Dizdaroglu M, Clarke ND (2000) Assignment of Enzyme Substrate Specificity by Principal Component Analysis of Aligned Protein Sequences: An Experimental Test Using DNA Glycosylase Homologs. Proteins 40(1): 98–105

46. Baxter-Koenigs AR, El Nesr G, Barrick D, Doug Barrick C, Jenkins T (2022) Singular value decomposition of protein sequences as a method to visualize sequence and residue space. Protein Sci. 31:e4422.

47. Qin C, Colwell LJ (2018) Power law tails in phylogenetic systems. Proc. Natl. Acad. Sci. U. S. A. 115:690–695.

48. Wheeler LC, Lim SA, Marqusee S, Harms MJ (2016) The thermostability and specificity of ancient proteins. Curr. Opin. Struct. Biol. 38:37–43.

49. Mitchell AL, Attwood TK, Babbitt PC, Blum M, Bork P, Bridge A, Brown SD, Chang HY, El-Gebali S, Fraser MI, et al. (2019) InterPro in 2019: Improving coverage, classification and access to protein sequence annotations. Nucleic Acids Res. 47:D351–D360.

50. Li W, Godzik A (2006) Cd-hit: A fast program for clustering and comparing large sets of protein or nucleotide sequences. Bioinformatics 22:1658–1659.

51. Katoh K, Standley DM (2013) MAFFT multiple sequence alignment software version 7: Improvements in performance and usability. Mol. Biol. Evol. 30:772–780.

52. Sievers F, Wilm A, Dineen D, Gibson TJ, Karplus K, Li W, Lopez R, McWilliam H, Remmert M, Söding J, et al. (2011) Fast, scalable generation of high-quality protein multiple sequence alignments using Clustal Omega. Mol. Syst. Biol. 7.

53. Stothard P (2000) The Sequence Manipulation Suite: JavaScript programs for analyzing and formatting protein and DNA sequences. Biotechniques 28:1102–1104.

54. Katayanagi K, Miyagawa M, Matsushima M, Ishikawa M, Kanaya S, Nakamura H, Ikehara M, Matsuzaki T, Morikawa K (1992) Structural details of ribonuclease H from Escherichia coli as refined to an atomic resolution. J. Mol. Biol. 223:1029–1052.

