## supplemental information for "The importance of input sequence set to consensus-derived proteins and their relationship to reconstructed ancestral proteins"

**Supplement**

The average tail length for the RNases H used to derive the phylogenetic tree is 6 ± 5 residues (Figure S1A). Therefore, we wondered if the unusual behavior for WholeCons* is a direct result of the lack of this C-terminal tail. When the ten-residue tail present in ecRNH is added to the C-terminus of WholeCons*, a full-length protein is not produced, and instead results in a truncated protein missing the C-terminal tail and part of the terminal E-helix (as determined by mass spectrometry). Thus, the addition of the ecRNH tail to WholeCons* may cause local unfolding events that result in a protease sensitive C-terminal region. The stability of this protein was not characterized.

We then relaxed the criteria for gaps, or missing residues to be included in the generation of consensus residues to create a consensus with tail residues. The consensus proteins discussed thus far were designed with the rule that a consensus residue is ‘-‘ (none) if a gap exists in greater than 50% of the sequences in the MSA at that position (see Methods). Raising this requirement to greater than 80% of the sequences results in more MSA positions having a consensus residue that is not ‘-‘. Using this modified approach on the set of 405 RNase H sequences produces a WholeCons* protein with an additional six-residue C-terminal tail that consists of the same residues as the start of the ecRNH tail (LEDTGY). The thermodynamics of this variant (WholeCons* with ConsTail) were comparable to WholeCons*, with a similar stability and *m*-value (Figure S1, Table S1), suggesting the lack of a C-terminal tail is not the cause of deviation from typical RNase H behavior.

The reconstructed ancestors of RNase H contain a C-terminal tail (see Figure 1). Therefore, we asked the converse question – does this tail contribute to the observed stability and cooperativity of AncCcons*? Removing the nine-residue tail on AncCcons* does result in a protein that is destabilized with a lower the *m*-value (Figure S1, Table S1), as expected since the C-terminal residues are also required for ecRNH stability.

Interestingly, there is a homolog in the same clade as ecRNH, from *S. enterica*, that has a tail that is only one residue. We expressed and characterized this homolog to see how it compares to other RNases H. This protein has an *m*-value even more depressed than WholeCons* and AncCcons* NoTail. Thus, a naturally occurring protein without a C-terminal tail exists in nature and is presumed to be functional for the organism. Together, these data indicate that the lack of a C-terminal tail does not seem to be the cause of the deviation from two-state behavior for WholeCons* and shows that some RNases H require this extension for function while others do not.

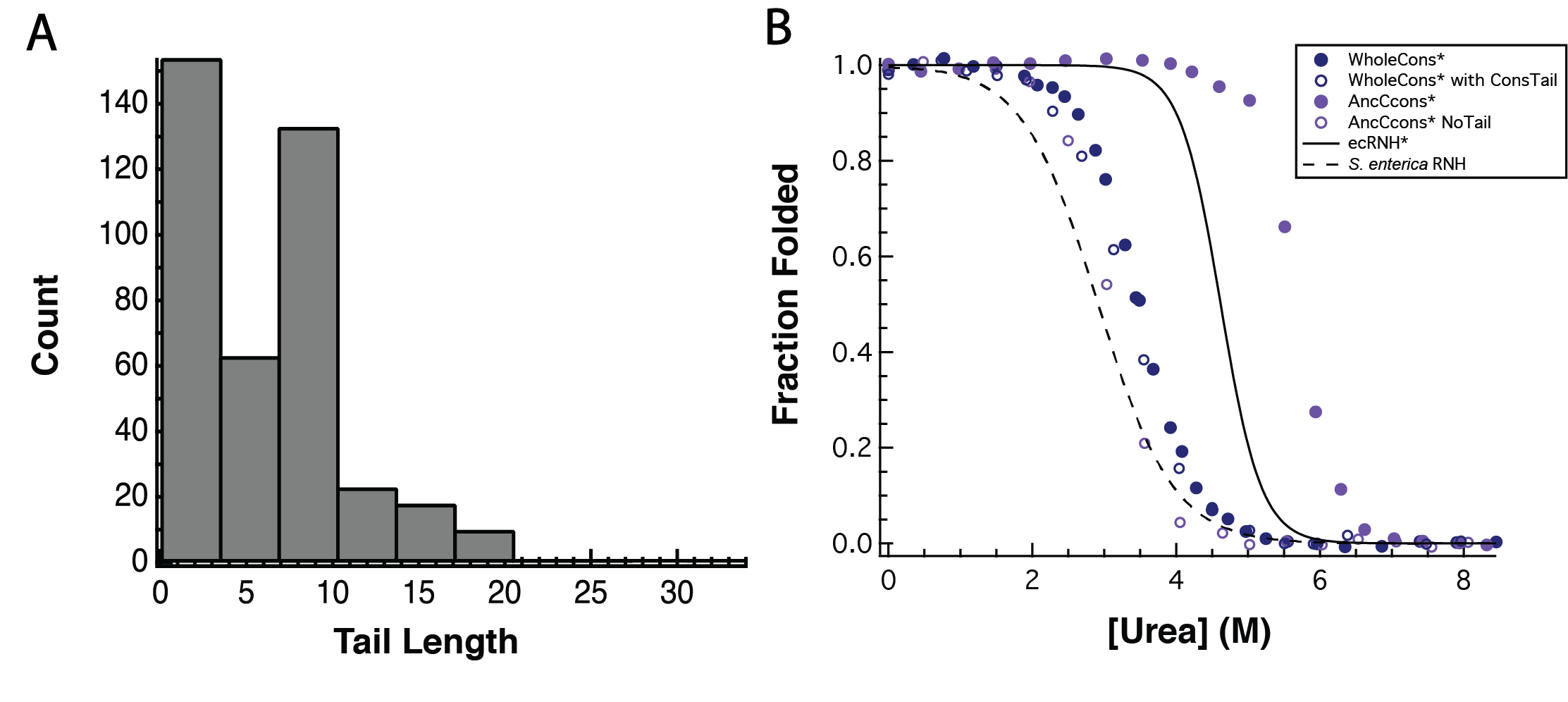

**Figure S1:** (**A**) Histogram of C-terminal tail lengths (# of residues C-terminal of residue 145 in ecRNH in the MSA) for the 405 RNase H homologs used to design WholeCons*. (**B**) Denaturant melts monitored by CD at 222 nm, pH 5.5, 25 °C for select RNases H with (designed WholeCons* with ConsTail, designed AncCcons*, extant ecRNH^24^) and without (designed WholeCons*, designed AncCcons* NoTail, extant S. enterica RNH) tails.

|  | Tm (°C) | ΔG_unf_ (kcal/mol) | *m*-value (kcal/mol/M) | Cm (M) |
| --- | --- | --- | --- | --- |
| ecRNH* | 68.0 ±0.5 | 9.7 ±0.4 | 2.1 ±0.09 | 4.62 ±0.04 |
| WholeCons* | 62.8 ±0.2 | 5.1 ±0.1 | 1.5 ±0.1 | 3.5 ±0.1 |
| WholeCons* with ConsTail | - | 4.6 ±0.2 | 1.4 ±0.04 | 3.4 ±0.03 |
| AncCcons* | 76.3 ±2.3 | 11.8 ±0.6 | 2.1 ±0.1 | 5.7 ±0.1 |
| AncCcons* No Tail | 58.5 ±0.4 | 5.3 | 1.7 | 3.1 |
| *S. enterica* RNH | 55.5 | 3.4 ±0.1 | 1.2 ±0.1 | 2.9 ±0.1 |

**Table S1:** Table of thermodynamic parameters from fits of Figure S1B (pH 5.5, 25 °C). Errors reported are standard deviations of replicate experiments. ecRNH* data from Raschke et al., Nat. Struct. Biol., 1999^18^ and S. enterica RNH T_m_ from Hart et al., PLoS Bio., 2014.
